# Genome-wide transcriptional responses of iron-starved *Chlamydia trachomatis* reveal prioritization of metabolic precursor synthesis over protein translation

**DOI:** 10.1101/217992

**Authors:** Amanda J. Brinkworth, Mark R. Wildung, Rey A. Carabeo

## Abstract

Iron is essential for growth and development of *Chlamydia*. Its long-term starvation in cultured mammalian cells leads to production of aberrant non-infectious chlamydial forms, also known as persistence. Immediate transcriptional responses to iron limitation have not been characterized, leaving a knowledge gap of how *Chlamydia* regulates its response to changes in iron availability. We used the fast-chelating agent 2,2’-Bipyridyl (BPDL) to homogeneously starve *Chlamydia trachomatis* serovar L2 of iron, starting at 6 or 12h post-infection. Immediate transcriptional responses were monitored after only 3 or 6h of BPDL-treatment, well before formation of aberrant *Chlamydia.* The first genome-wide transcriptional response of *C. trachomatis* to iron-starvation was subsequently determined utilizing RNA-sequencing. Only 7% and 8% of the genome was differentially expressed in response to iron-starvation at early and mid-stages of development, respectively. Biological pathway analysis revealed an overarching theme. Synthesis of macromolecular precursors (deoxynucleotides, amino acids, charged tRNAs, and acetyl-coA) was up-regulated, while energy-expensive processes (ABC transport and translation) were down-regulated. A large fraction of differentially down-regulated genes are involved in translation, including ribosome assembly, initiation and termination factors, which could be analogous to the translation down-regulation triggered by stress in other prokaryotes during stringent responses. Additionally, transcriptional up-regulation of DNA repair, oxidative stress, and tryptophan salvage genes reveals a possible coordination of responses to multiple antimicrobial and immunological insults. These responses of replicative-phase *Chlamydia* to iron-starvation indicate a prioritization of survival over replication, enabling the pathogen to “stock the pantry” with ingredients needed for rapid growth once optimal iron levels are restored.

**IMPORTANCE:** By utilizing an experimental approach that monitors the immediate global response of *Chlamydia trachomatis* to iron-starvation, clues to long-standing questions in *Chlamydia* biology are revealed, including how *Chlamydia* adapts to this stress. We determined that this pathogen initiates a transcriptional program that prioritizes replenishment of nutrient stores over replication, possibly in preparation for rapid growth once optimal iron levels are restored. Transcription of genes for biosynthesis of metabolic precursors was generally up-regulated, while those involved in multiple steps of translation were down-regulated. We also observed an increase in transcription of genes involved in DNA repair and neutralizing oxidative stress, indicating that *Chlamydia* employs an “all-or-nothing” strategy. Its small genome limits its ability to tailor a specific response to a particular stress. Therefore, the “all-or-nothing” strategy may be the most efficient way of surviving within the host, where the pathogen likely encounters multiple simultaneous immunological and nutritional insults.

## INTRODUCTION

The sexually transmitted bacterium *Chlamydia trachomatis* infects the mucosal epithelium of the endocervix, urethra, and anogenital tract. These infections usually resolve spontaneously, and most are asymptomatic and thus underreported. Over 1.5 million cases of *C. trachomatis* genital infections were reported in the US in 2015 alone (1). As many as 17% of females infected with *C. trachomatis* develop long-term infections in the genital tract, which can result in serious complications such as pelvic inflammatory disease (PID), fallopian-tube scarring, and ectopic pregnancy, all of which are major risk factors for tubal factor infertility (TFI) (2). Rectal infections with LGV serovars of *C. trachomatis* can be invasive, and if untreated can lead to complications such as proctocolitis, inguinal adenopathy, reactive arthropathy and colorectal ulcers (3). In some patients, infection persists even after antibiotic treatment (4, 5). The ability of *C. trachomatis* to survive long-term in some individuals despite host immunity and antibiotic treatment is not well understood, and may be associated with *Chlamydia*’s ability to become persistent (6). While aberrant chlamydial forms have been identified in cervical samples, the clinical relevance of this phenomenon is not well understood (6, 7).

Chlamydiae are obligate intracellular Gram-negative bacteria that undergo a biphasic developmental cycle that includes both non-replicative and replicative forms (8). Infection begins when the small, metabolically quiescent chlamydial elementary body (EB) binds to mucosal epithelial cells and translocates virulence factors that induce its endocytic uptake. Within 2 hours of entry, the EB will differentiate into its replicative form, the reticulate body (RB). Continued secretion of effectors leads to modification of the endocytic vesicle such that it avoids fusion with the lysosome and enables capture of nutrient-rich vesicles. This unique intracellular niche, called the inclusion, continues to expand as RBs replicate. In response to unknown signals around 24h post-infection, RBs will then differentiate into infectious EBs, followed by EB release 36-72 hours post-infection (8). Under exposure to certain stress conditions in cell culture (e.g. penicillin, interferon-gamma (IFN-g), iron-depletion or tryptophan-depletion) RBs will not differentiate into EBs, but instead enter into a state of persistence, characterized by aberrant, enlarged morphology (9–13). Persistent *Chlamydia* are resistant to both antibiotics and host immunity mechanisms, and can recover from this state upon removal of stress or addition of missing nutrients (14-17).

Chlamydiae have undergone reductive evolution as they have adapted to intracellular growth in mammalian cells, discarding metabolic genes responsible for synthesizing factors that could be acquired from the host (18). The core genome of *C. trachomatis* serovar L2 encodes only 889 open-reading frames, making *Chlamydia* dependent on its host for lipids, nucleotides, amino acids, and metal cofactors (18). Exposure of *Chlamydia-*infected cells to immune mediators, such as IFN-g, reduces the availability of these factors and results in reduced RB division and differentiation (12, 14). IFN-g induces intracellular depletion of tryptophan by increasing levels of indolamine 2,3-dioxygenase, which is responsible for catabolizing tryptophan into kynurenines, which cannot be utilized in tryptophan metabolism (19).

Induction of inflammatory cytokines such as IFN-g and IL-6 in response to chlamydial infection likely causes sequestration of free iron by the mononuclear phagocytic system, which includes both cellular and systemic regulatory pathways (20–27). Readers are referred to two comprehensive reviews of the coordinated regulation of iron homeostasis by systemic and cellular mechanisms (26, 28). In the context of *Chlamydia* infection of the genital epithelium, iron availability in infected cells is likely limited by down-regulation of transferrin receptor and upregulation of the iron-storage factor, ferritin (29). Iron-levels in the female genital tract can also fluctuate throughout the menstrual cycle, in part due to hormone-induced expression of lactoferrin (30, 31). Iron is essential for growth and development of *Chlamydia*, and its acquisition and accumulation must be carefully regulated. In mammals, the readily usable ferrous iron (Fe^2+^) is tied up in molecular complexes, limiting their interaction with hydrogen peroxide to form damaging hydroxyl radicals through the Fenton reaction (32). Eukaryotic stores of ferric iron (Fe^3+^) are strictly regulated to restrict access by pathogenic bacteria (33). Extracellular bacteria such as *Pseudomonas* and *Yersinia* utilize multiple redundant iron-binding molecules called siderophores that compete with mammalian transferrin for ferrous iron (34, 35). Intracellular bacteria, such as *Mycobacterium*, *Francisella* and *Chlamydia*, can obtain iron by subverting host vesicles that contain holo-transferrin bound to transferrin receptor (36-39). Using a combination of endocytic markers and chemical inhibitors, our laboratory discovered that *Chlamydia* specifically recruits transferrin-containing vesicles from the slow-recycling endocytic pathway (38). Once delivered into the inclusion, iron is likely imported into bacteria through an ABC transporter system, encoded by *ytgABCD*, which is the only known iron acquisition system in *Chlamydia* species (40-42). The C-terminus of the YtgC permease, referred to as YtgR, is homologous to the *Cornyebacterium* repressor DtxR, and has been recently characterized as an iron-dependent repressor of the *ytgABCD* iron-acquisition operon (42). A recent review highlights the differences between iron-acquisition strategies of *Chlamydia* with other intracellular bacteria (27).

Conversion to the aberrant phenotype in response to iron-starvation reduces the infectious potential of *Chlamydia*, since only a portion of RBs recover from stress and complete development into infectious EBs once iron is added back into the media (43). Previous studies have characterized aberrant *C. trachomatis* after long-term treatment with the iron-chelator deferoxamine, and have detected increased expression of the iron-binding protein YtgA, indicating its role in iron-uptake (41, 44, 45). However, these studies added deferoxamine at the time of infection, and did not monitor transcriptional or proteomic patterns until ≥24 hours post-infection, making it difficult to determine if this up-regulation is part of the initial response to iron starvation. Immediate genome-wide transcriptional responses to iron limitation have not yet been characterized in detail, leaving a gap in the current knowledge of how *Chlamydia* regulates its response to changes in iron availability. We utilize the chelator 2,2-Bipyridyl (BPDL), which can quickly and efficiently chelate free iron from both bacterial and mammalian cells, to induce an immediate transcriptional response to iron-starvation by *C. trachomatis* (43, 46–48).

This study provides the first global profile of the *C. trachomatis* transcriptional response to iron starvation. Our short-term, effective treatment regimen in combination with deep RNA-sequencing reveals the immediate response of *Chlamydia* to iron-starvation in the logarithmic phase of growth when the bacteria are in the RB form. Here we utilize this dataset to map the specific biological pathways altered in response to iron-starvation. Taken together our results provide important clues to how *Chlamydia* survives iron limitation. Accumulation of metabolite precursors is prioritized over macromolecular biosynthesis. In addition, the transcriptional induction of genes involved in adaptation to other stress factors, e.g. oxidative stress and amino acid starvation, points to the inability of *Chlamydia* to tailor its transcriptional response to a specific stress. Lastly, the global transcriptomic profile of iron-starved *Chlamydia* provided valuable insights into how the biphasic developmental cycle might irreversibly switch to persistence.

## RESULTS

### Treatment optimization to detect the immediate chlamydial response to iron-starvation

The bivalent chelator 2,2-Bipyridyl (BPDL) has been shown to deplete both ferrous and ferric iron from *Chlamydia*-infected cells during long-term treatment, and it induces the development of aberrant forms more consistently and homogenously than the previously used ferric iron chelator, deferoxamine (43). Here, we determined the optimal duration of BPDL treatment to induce iron-responsive transcription without inducing morphological abnormalities in *Chlamydia*. We chose to begin starvation during mid-cycle development (12h p.i.) instead of at the beginning of infection for two reasons: 1) to test the response of actively replicating *Chlamydia* that are able to maximally respond to stress, and 2) to ensure that both treated and mock-treated *Chlamydia* remain in the same stage of development (RB). We monitored chlamydial morphology, growth, and transcriptional responses after 3, 6, and 12h of BPDL treatment (Figure 1A). Indirect immunofluorescent confocal microscopy revealed similar morphology between mock-treated and BPDL-treated forms for up to 12h of BPDL-treatment (Figure 1B). Interestingly, observation of BPDL-treated cultures by light microscopy revealed an obvious decrease in Brownian movement within inclusions after 6 or more hours of treatment (data not shown). This observation is consistent with our findings that chlamydial growth is reduced compared to mock-treated after only 6h of BPDL-treatment, as determined by quantitative PCR of chlamydial genomes (Figure 1C).

**Figure 1.**
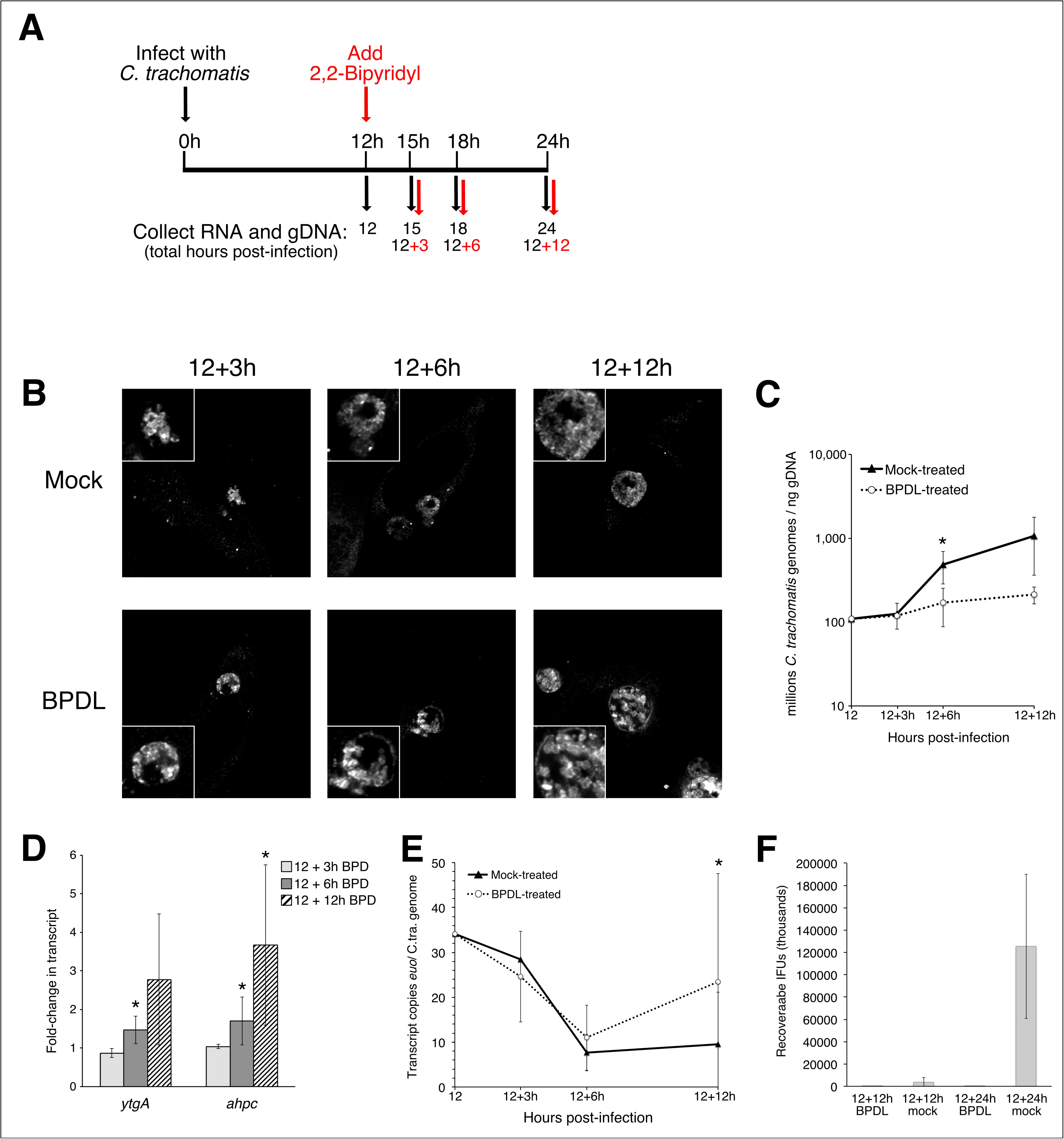
Optimization of 2,2-Bipyrdyl (BPDL) treatment to induce iron-responsiveness in the absence of persistence. Timeline of BPDL-treatment A) starting at 12h p.i., BPDL was supplemented to culture media for 3, 6, or 12h. Mock-treated and BPDL-treated samples were tested for changes to B) morphology by confocal microscopy, C) growth by qPCR, D) iron-responsive transcription (*ytgA, ahpC*) D), and transcription of the developmental marker, *euo*, by RT-qPCR. Significant changes with a p-value ≤ 0.05 in a one-tailed Student’s t-test are indicated with an asterisk, and are based on 3 biological replicates for the growth curve and 4 biological replicates for RT-qPCR.

We monitored the transcriptional response of the known iron-responsive genes *ytgA* and *ahpC* by reverse transcriptase quantitative polymerase chain reaction (RT-qPCR) to validate the iron starvation protocol (40, 41, 43, 49). Elevated transcription of both iron-starvation markers was detected after only 6h BPDL-treatment, compared to mock-treated (1.5-fold and 1.7-fold). Maximum differences in transcription of both markers were detected after 12h of BPDL treatment (2.8-fold and 3.7-fold; Figure 1D). In the same experiment, we also monitored the transcriptional profile of the early gene, *euo*, which decreases during late stages of normal development. Multiple persistence models have demonstrated dysregulated of *euo* transcription, with high levels of *euo* mRNA detected late in development under persistence-inducing conditions (15, 43, 45). After 12h BPDL treatment, we observed that *euo* transcript levels remained elevated relative to the mock-treated control, indicating dysregulated transcription or a possible delay in development (Figure 1E, top). This delay in development during longer BPDL-treatment is supported by the lack of recoverable inclusion-forming units (IFUs) detected after 12 or 24h BPDL-treatment compared to mock-treated controls (Figure 1F), indicating a possible lack of RB-to-EB differentiation. Because 6h BPDL-treatment is sufficient to induce iron-responsive transcription without inducing the morphology and transcriptional pattern associated with persistence, we chose it as the optimal duration of iron-starvation for our genome-wide transcriptional studies. We also included 3h BPDL-treatment to detect the earliest possible response to iron-starvation prior to BPDL-induced changes in growth.

### Global transcriptional response of *C. trachomatis* to iron-starvation during mid-cycle development

The primary global response of *C. trachomatis* to mid-cycle iron starvation was determined by RNA-sequencing (RNA-seq). We utilized an Ion Proton for sequencing, which allowed for easy and rapid scaling of timepoints based on observed yield of mapped reads. This approach is relevant to the study of *Chlamydia* transcription because chlamydial mRNA represents a small proportion of the total RNA at the time points analyzed, even after significant enrichment steps. For mid-cycle iron starvation studies, we aimed for greater than 10x coverage of 100% of the *C. trachomatis* genome, with a minimum of 3 biological replicates per sample. The sequencing reads were trimmed to exclude adaptor sequences and polyclonal reads, followed by exclusion of reads less than 30 nucleotides in length. The remaining reads were aligned to the *C. trachomatis* genome and plasmid, with 2 to 23% of trimmed reads mapping. Average read lengths varied from 92 to 134 nucleotides, requiring an average of 108,837 mapped reads to reach our coverage goal. A summary of the read and mapping statistics for all of our samples can be found in Supplementary Table 1. Alignments, comparisons, and normalization of aligned reads were done with CLC Genomics Workbench version 9.0 according to default settings. All mid-cycle conditions (12h untreated, 12+3h BPDL, 12+3h mock, 12h+6h BPDL, 12+6h mock) were compared using the CLC Genomics experiment tool, normalized by quantile scaling and analyzed for differential gene expression using EdgeR statistical analysis with false-discovery rate (FDR) calculation. Because we included ribosomal rRNA, eukaryotic mRNA, and small RNA (<100 nt) depletion steps when preparing chlamydial mRNA for RNA-seq, we also excluded tRNAs, rRNAs, and genes with < 10 mean reads in a sample group prior to normalization and analysis.

The genome-wide profile of mock-treated and BPDL-treated gene expression during mid-cycle development (12h to 18h post-infection) is displayed as a heatmap of normalized expression values (Figure 2A). The raw and normalized data for these individual replicates can be found in Supplemental Table 4. Mock-treated (left) and BPDL-treated (right) profiles are remarkably similar across all genes that significantly change during normal mid-cycle development of *Chlamydia* (based on comparisons 12h vs 15h, 15h vs 18h, or 12h vs 18h, p-value ≤ 0.01). The annotated expression heatmap and EdgeR comparisons for normal growth can be found in Supplemental Figure 1 and Supplemental Table 2, respectively. The entire RNA-seq dataset of normal development can be found in Supplemental Table 3. The similarity between global expression profiles indicates that *Chlamydia’s* normal development is not dysregulated after only 3h or 6h of BPDL-treatment. However, EdgeR analysis of BPDL-treated cultures compared to mock-treated samples (at equivalent time points post-infection) revealed 8% (76/889) of the genome was differentially expressed after 3h BPDL–treatment and 1% (12/889) was differentially expressed after 6h BPDL-treatment. Genes that were differentially expressed with a maximum p-value of 0.01 are displayed in a heatmap of fold-change differences between BPDL-treated and mock-treated samples (Figure 2B). Examples of decreased transcription after 3h and 6h BPDL-treatment include the ribosomal subunit genes *rpsO* and *rpsT*, and the type III secretion genes *copB* and *scc2*, respectively. Transcription of the tryptophan salvage pathway operon, *trpBA*, and the ribonucleotide reductase operon, *nrdAB*, was significantly increased after both 3 and 6h of treatment. Iron-responsive genes that were differentially expressed with a p-value ≤ 0.01 after 3 or 6h of BPDL-treatment are listed in Table 1 and Table 2, respectively. The fully annotated heatmap can be found in Supplemental Figure 2, and the full set of RNA-sequencing results for mid-cycle iron starvation can be found in Supplemental Table 4.

**Figure 2.**
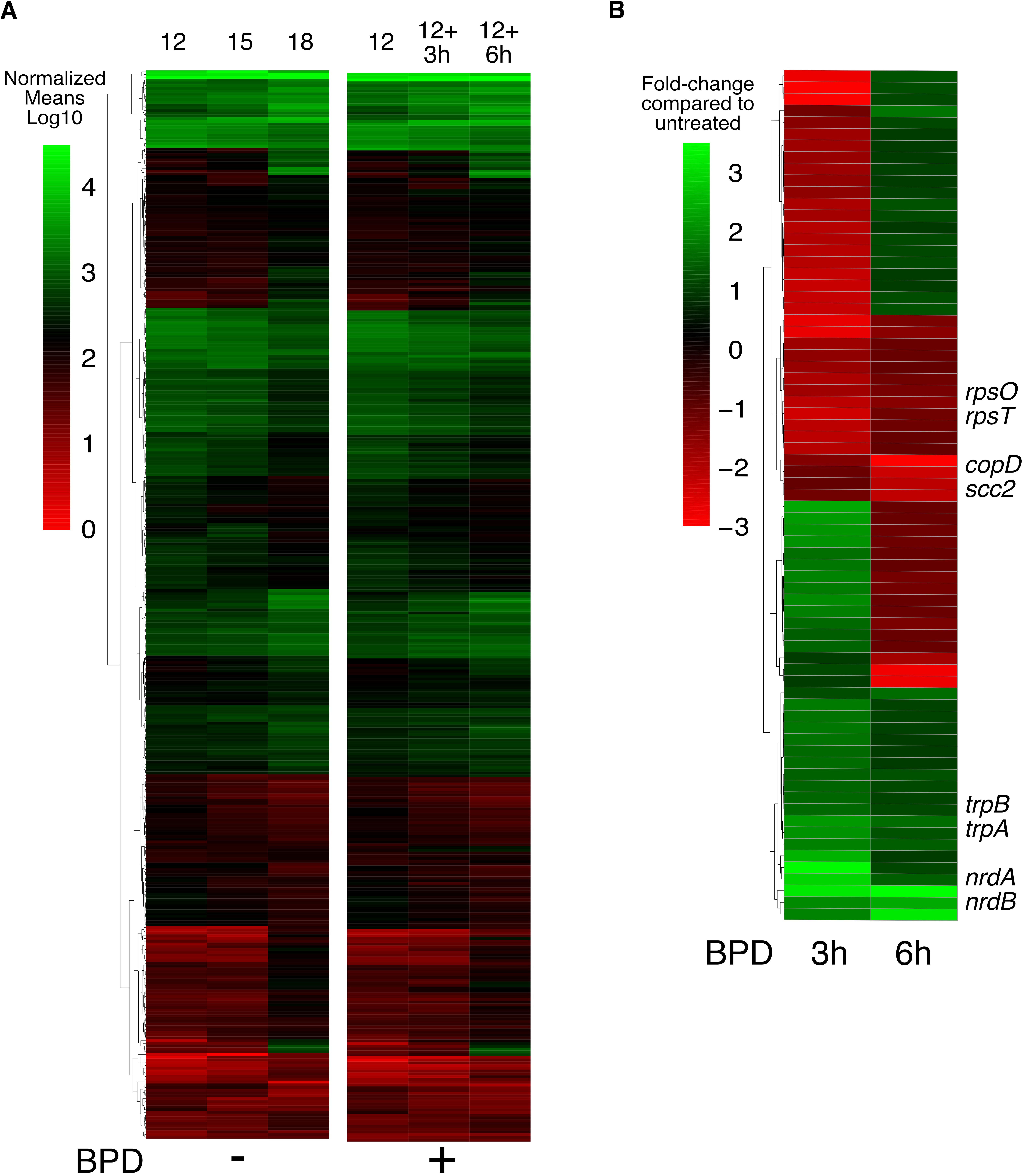
Global and differential gene expression of the mid-cycle response to iron-starvation. The global response of *C. trachomatis* to iron starvation was detected by RNA-sequencing and reads were aligned to the genome and plasmid. A) The untreated expression profile is displayed for all genes that change significantly (p-value ≤ 0.01) during mid-cycle development as a heatmap of log10 transformed normalized expression means (left). Expression across the same genes are displayed for BPDL-treated samples (right). The highest and lowest expression values are displayed in green and red, respectively. B) Genes that are significantly changed in response to iron-starvation, with a p-value ≤ 0.01, are displayed as a heatmap of fold-changes of BPDL-treated compared to mock-treated equivalent samples. The most highly up-regulated and down-regulated transcripts are displayed in green and red, respectively.

**Table 1.**
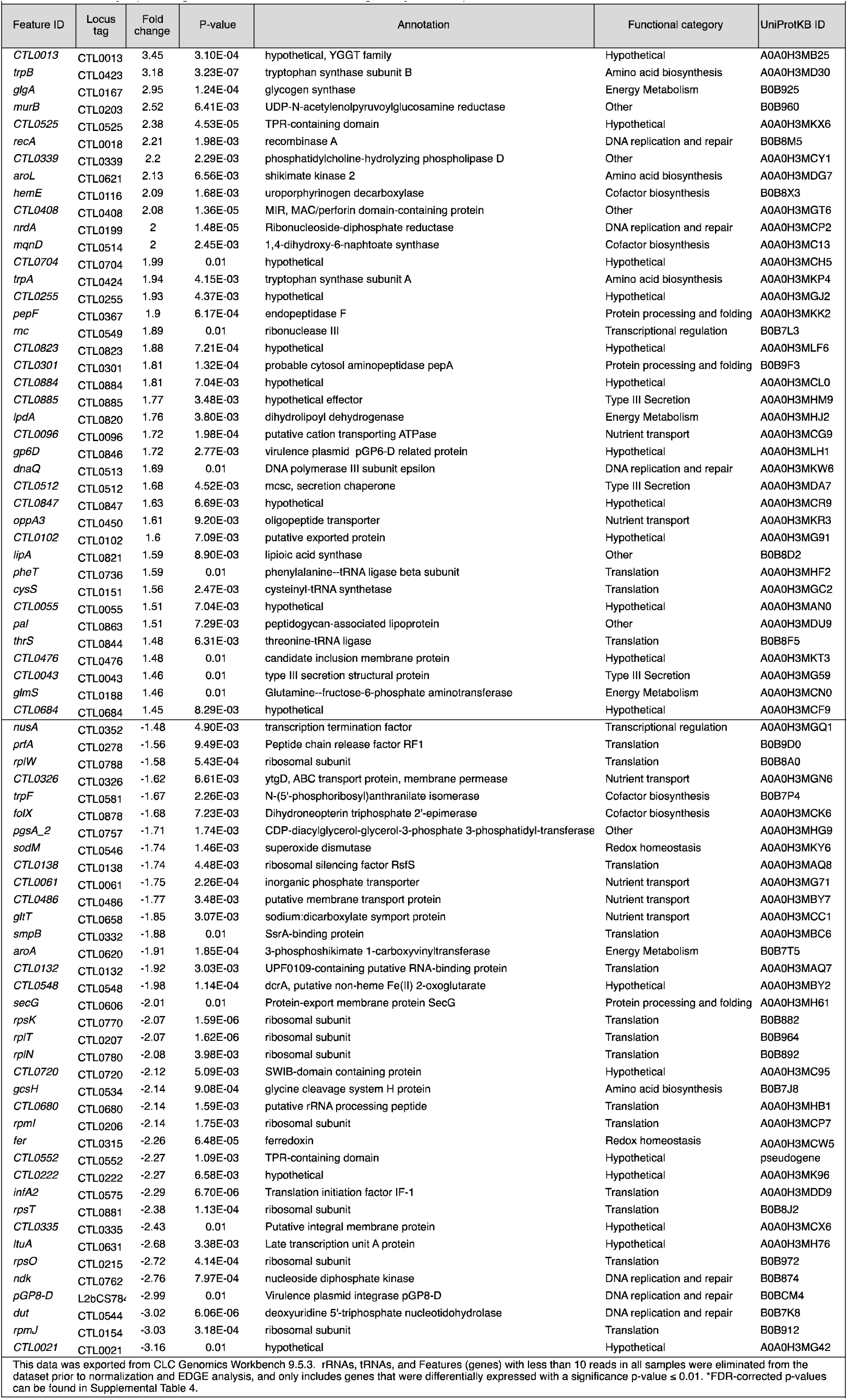
Differentially expressed genes after 3h BPD treatment during mid-cycle development.

**Table 2.** Differentially expressed genes after 6h BPD treatment during mid-cycle development.

|  |  |  |  |  |  |  |
| --- | --- | --- | --- | --- | --- | --- |
| Feature ID | Locus Tag | P-value | Fold change | Annotation | Functional category | UniProtKB ID |
| *trpB* | CTL0423 | 2.61E-07 | 3.5 | tryptophan synthase subunit B | Amino acid biosynthesis | A0A0H3MD30 |
| *trpA* | CTL0424 | 9.26E-06 | 3.21 | tryptophan synthase subunit A | Amino acid biosynthesis | A0A0H3MKP4 |
| *nrdA* | CTL0199 | 3.50E-06 | 2.39 | ribonucleoside-diphosphate reductase | DNA replication and repair | A0A0H3MCP2 |
| *CTL0071* | CTL0071 | 6.73E-03 | 1.73 | hypothetical | Hypothetical | A0A0H3MCG5 |
| *nrdB* | CTL0200 | 9.34E-03 | 1.63 | ribonucleoside-diphosphate reductase | DNA replication and repair | A0A0H3MK81 |
| *CTL0619* | CTL0619 | 1.68E-03 | -1.83 | hypothetical integral membrane protein | Hypothetical | A0A0H3MH71 |
| *copD* | CTL0842 | 3.17E-03 | -2.13 | type III secretion system protein | Type III Secretion | A0A0H3MHK7 |
| *scc2* | CTL0839 | 1.37E-03 | -2.18 | type III secretion system chaperone | Type III Secretion | A0A0H3MLG7 |
| *tsp* | CTL0700 | 1.28E-03 | -2.34 | tail-specific protease | Protein processing and folding | A0A0H3MDM0 |
| *CTL0185* | CTL0185 | 6.85E-03 | -2.76 | hypothetical membrane protein | Hypothetical | A0A0H3MAV2 |
| *copB* | CTL0841 | 6.79E-05 | -2.81 | type III secretion system membrane protein | Type III Secretion | A0A0H3MCF0 |
| *CTL0840* | CTL0840 | 2.15E-03 | -2.97 | hypothetical | Hypothetical | A0A0H3MCQ4 |
| This data was exported from CLC Genomics Workbench 9.5.3. rRNAs, tRNAs, and Features (genes) with less than 10 reads in all samples were eliminated from the dataset prior to normalization and EDGE analysis. This data only includes genes that were differentially expressed with a significance p-value $\leq 0.01$. *FDR-corrected p-values can be found in Supplemental Table 4. | | | | | | |

### Functional categorization of differentially expressed genes during mid-cycle response to iron-starvation

Annotations and functional categories of differentially expressed genes during mid-cycle iron-starvation were retrieved from UniProt and are listed in Table 1 and Table 2. Genes differentially expressed, with a minimum p-value of 0.01, after only 3h of BPDL-treatment are grouped by functional category of induced and reduced transcripts (Table 1, Figure 3) (50). Of the 39 genes significantly induced after only 3h iron-starvation, representing 4% of the genome, five categories were equally represented with 3 genes each: energy metabolism (*glgA, lpdA, glmS*), amino acid biosynthesis (*trpA, trpB, aroL*), DNA replication and repair (*nrdA, recA, dnaQ*), type III secretion (*mcsC, ctl0085, ctl0043*), and translation (*pheT, cysS, thrS*). Of the 37 genes that were significantly reduced in response to 3h BPDL-treatment, representing 4% of the genome, the majority of these (39%) are associated with translation (*prfA, rplW, rsfS, smpB, ctl0132, rpsK, rplT, rplN, ctl0680, rpmI, rmpJ, infa2, rpsT, rpsO*), and 11% are associated with nutrient transport (*ctl0061, ctl0485, gltT*, and *ytgD*). Transcript levels of only 5 genes were significantly increased after 6h BPDL-treatment (*trpB, trpA, nrdA, ctl0071*, and *nrdB*) while transcript levels of 7 genes were decreased (*ctl0185, ctl0619, tsp*, and the entire *scc2-ctl0840-cobB-copD* operon) (Table 2).

**Figure 3.**
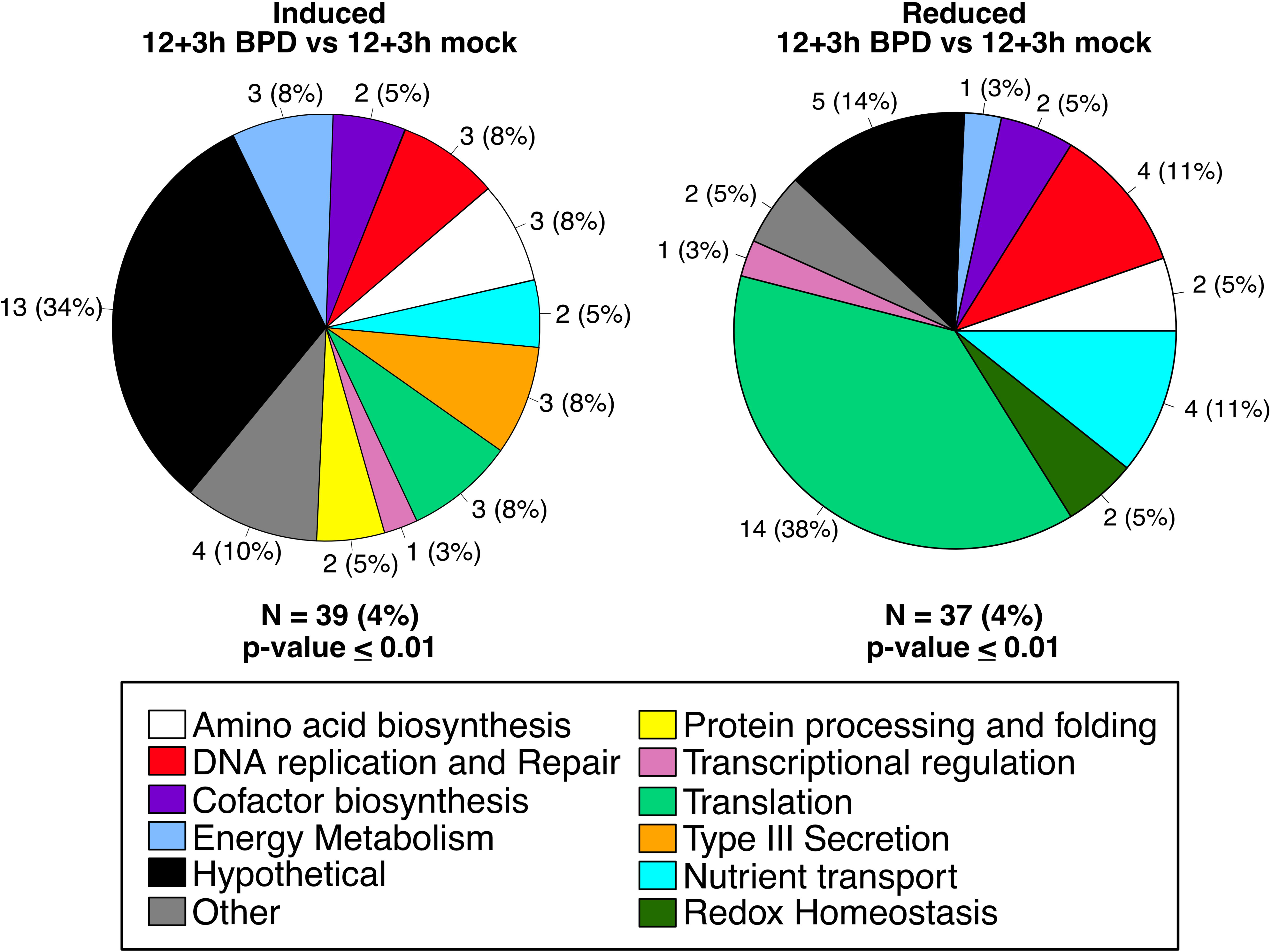
Functional categorization of mid-cycle response to iron-starvation. Transcripts that were significantly up-regulated (left) or down-regulated (right) after 3h BPDL treatment, starting at 12h p.i., are organized in pie charts by their functional categories. Adjacent to each pie slice is the number of genes in that category, and in parenthesis is the percentage of differentially expressed genes in the category. N=number of differentially expressed genes, and the percentage of the total genome that is represented.

To independently confirm the mid-cycle response detected by RNA-sequencing, we utilized RT-qPCR. Increased transcription in response to iron-starvation was confirmed for all of the transcripts tested by RT-qPCR, with the exception of *recA* (Figure 4A). None of the tested down-regulated genes were significantly reduced compared to controls by RT-qPCR, likely due to the fact that the genes tested were very low in abundance at the timepoints tested (Figure 4B).

**Figure 4.**
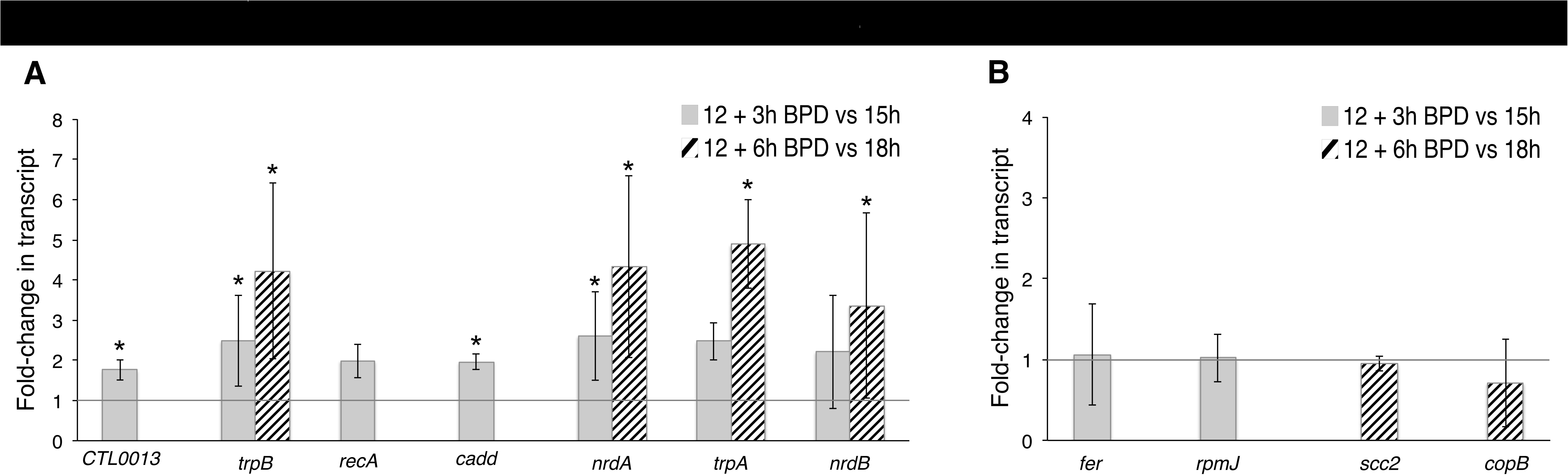
Confirmation of the mid-cycle response to iron starvation by RT-qPCR. Differentially expressed transcripts detected by RNA-sequencing were confirmed by RT-qPCR. A) Up-regulated or B) down-regulated transcription is indicated as fold-changes in transcripts after 3h BPD-treatment (solid gray bar) or 6h BPDL-treatment (striped bar) compared to mock-treated samples at equivalent timepoints post-infection. An asterisk indicates that the fold-change is significantly changed, with a p-value ≤ 0.05. Statistical analysis was done with a one-tailed Student’s t-test, based on at least 3 biological replicates.

### Functional categorization of differentially expressed genes during early-cycle response to iron-starvation

*Chlamydia* infections of the genital tract are asynchronous. Thus *Chlamydia* could be exposed to host-induced stress at any point in the developmental cycle. For this reason, we extended our analysis of the immediate response to iron starvation to an earlier point in the developmental cycle. *Chlamydia*-infected cells were treated with BPDL starting at 6h post-infection, which is after the initial EB-to-RB differentiation and at the beginning of the logarithmic growth phase. RNA and genomic DNA were collected at 9h post-infection for both treated and mock-treated samples. RNA-seq and alignments were performed as described above. A summary of mapped reads and coverage can be found in Supplemental Table 1.

Genes differentially expressed during the early cycle response (6+3h BPDL vs 6+3h mock-treated), with a maximum p-value of 0.01, are grouped by functional categories of induced and reduced transcripts (Figure 5). The full set of differentially expressed genes and their annotations can be found in Table 3. Similar to results of the mid-cycle response, transcription of 4% of the genome was induced, including genes involved in DNA replication and repair (*nrdA, nrdB, mutS, dnaQ*, and *recA*), amino acid biosynthesis (*trpB, trpA*, *aspC_1*, and *glyA*), and translation (*ctl0111*, *trpS*, *thrS*, and *aspS*) during the early-cycle response to iron starvation. Uniquely, genes involved in redox homeostasis (*pdi*, *ahpC* and *sodM*) were also up-regulated in response to iron-starvation starting at 6h post-infection but not during the mid-cycle response. Of the 23 genes with reduced transcription during the early-cycle response to iron-starvation (3% of the genome), 17% are associated with translation (*rplW, prfA, rplC, ftsY*), and 13% with DNA replication and repair (*pGP8D*, *amn*, and *dnaX_1*).

**Figure 5.**
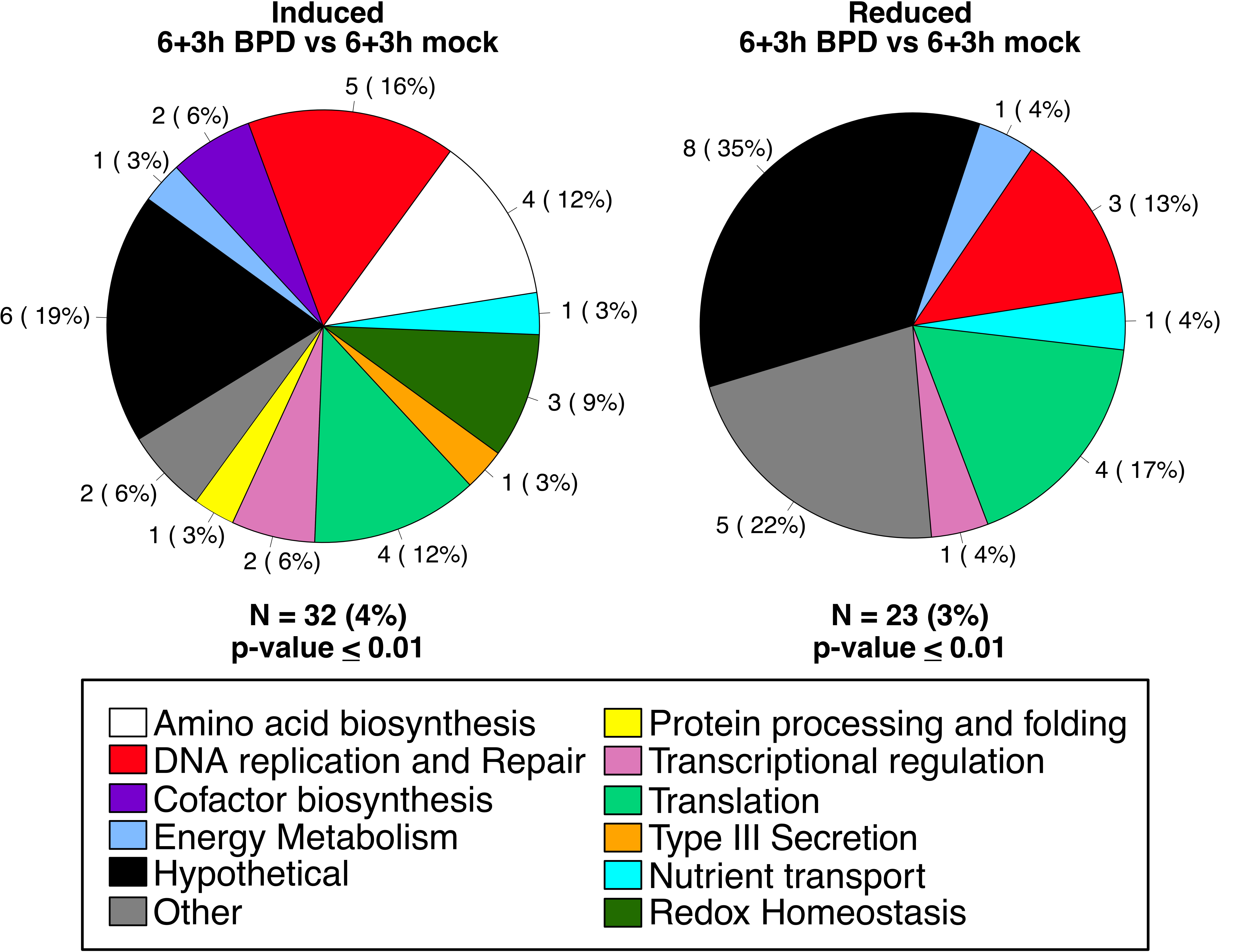
Functional categorization of the early-cycle response to iron-starvation. Transcripts that were significantly up-regulated (left) or down-regulated (right) after 3h of BPDL treatment, starting at 6h p.i., are organized in pie charts by their functional categories. Adjacent to each pie slice is the number of genes in that category, and in parenthesis is the percentage of differentially expressed genes in the category that make up the pie. N=number of differentially expressed genes, and the percentage of the total genome represented.

**Table 3.**
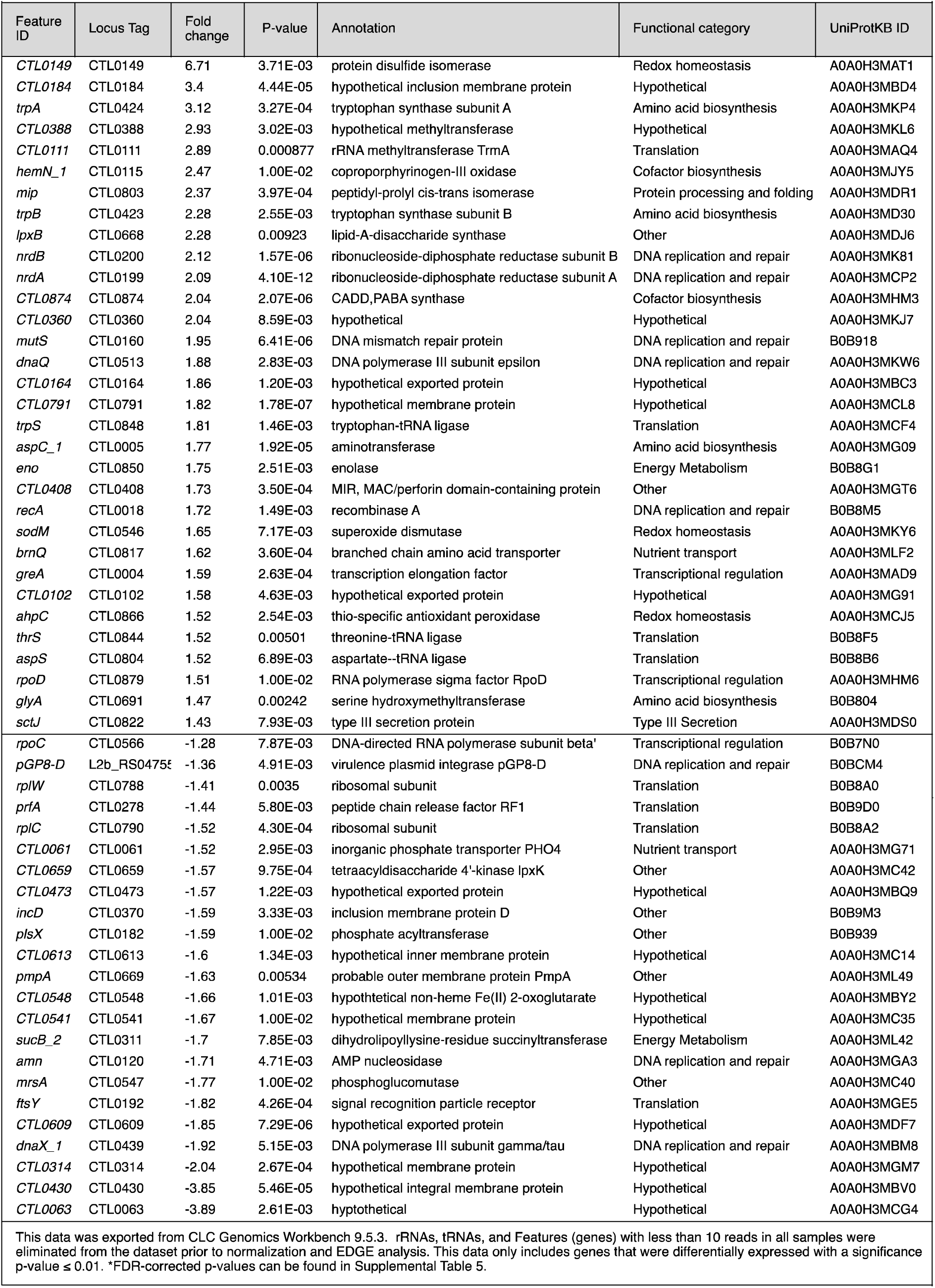
Differentially expressed genes after 3h BPD treatment during mid-cycle development.

| Feature ID | Locus Tag | Fold change | P-value | Annotation | Functional category | UniProtKB ID |
| --- | --- | --- | --- | --- | --- | --- |
| *CTL0149* | CTL0149 | 6.71 | 3.71E-03 | protein disulfide isomerase | Redox homeostasis | A0A0H3MAT1 |
| *CTL0184* | CTL0184 | 3.4 | 4.44E-05 | hypothetical inclusion membrane protein | Hypothetical | A0A0H3MBD4 |
| *trpA* | CTL0424 | 3.12 | 3.27E-04 | tryptophan synthase subunit A | Amino acid biosynthesis | A0A0H3MKP4 |
| *CTL0388* | CTL0388 | 2.93 | 3.02E-03 | hypothetical methyltransferase | Hypothetical | A0A0H3MKL6 |
| *CTL0111* | CTL0111 | 2.89 | 0.000877 | rRNA methyltransferase TrmA | Translation | A0A0H3MAQ4 |
| *hemN_1* | CTL0115 | 2.47 | 1.00E-02 | coproporphyrinogen-III oxidase | Cofactor biosynthesis | A0A0H3MJY5 |
| *mip* | CTL0803 | 2.37 | 3.97E-04 | peptidyl-prolyl cis-trans isomerase | Protein processing and folding | A0A0H3MDR1 |
| *trpB* | CTL0423 | 2.28 | 2.55E-03 | tryptophan synthase subunit B | Amino acid biosynthesis | A0A0H3MD30 |
| *lpxB* | CTL0668 | 2.28 | 0.00923 | lipid-A-disaccharide synthase | Other | A0A0H3MDJ6 |
| *nrdB* | CTL0200 | 2.12 | 1.57E-06 | ribonucleoside-diphosphate reductase subunit B | DNA replication and repair | A0A0H3MK81 |
| *nrDA* | CTL0199 | 2.09 | 4.10E-12 | ribonucleoside-diphosphate reductase subunit A | DNA replication and repair | A0A0H3MCP2 |
| *CTL0874* | CTL0874 | 2.04 | 2.07E-06 | CADD, PABA synthase | Cofactor biosynthesis | A0A0H3MHM3 |
| *CTL0360* | CTL0360 | 2.04 | 8.59E-03 | hypothetical | Hypothetical | A0A0H3MKJ7 |
| *mutS* | CTL0160 | 1.95 | 6.41E-06 | DNA mismatch repair protein | DNA replication and repair | B0B918 |
| *dnaQ* | CTL0513 | 1.88 | 2.83E-03 | DNA polymerase III subunit epsilon | DNA replication and repair | A0A0H3MKW6 |
| *CTL0164* | CTL0164 | 1.86 | 1.20E-03 | hypothetical exported protein | Hypothetical | A0A0H3MBC3 |
| *CTL0791* | CTL0791 | 1.82 | 1.78E-07 | hypothetical membrane protein | Hypothetical | A0A0H3MCL8 |
| *trpS* | CTL0848 | 1.81 | 1.46E-03 | tryptophan-tRNA ligase | Translation | A0A0H3MCF4 |
| *aspC_1* | CTL0005 | 1.77 | 1.92E-05 | aminotransferase | Amino acid biosynthesis | A0A0H3MG09 |
| *eno* | CTL0850 | 1.75 | 2.51E-03 | enolase | Energy Metabolism | B0B8G1 |
| *CTL0408* | CTL0408 | 1.73 | 3.50E-04 | MIR, MAC/perforin domain-containing protein | Other | A0A0H3MGT6 |
| *recA* | CTL0018 | 1.72 | 1.49E-03 | recombinase A | DNA replication and repair | B0B8M5 |
| *sodM* | CTL0546 | 1.65 | 7.17E-03 | superoxide dismutase | Redox homeostasis | A0A0H3MKY6 |
| *brnQ* | CTL0817 | 1.62 | 3.60E-04 | branched chain amino acid transporter | Nutrient transport | A0A0H3MLF2 |
| *greA* | CTL0004 | 1.59 | 2.63E-04 | transcription elongation factor | Transcriptional regulation | A0A0H3MAD9 |
| *CTL0102* | CTL0102 | 1.58 | 4.63E-03 | hypothetical exported protein | Hypothetical | A0A0H3MG91 |
| *ahpC* | CTL0866 | 1.52 | 2.54E-03 | thio-specific antioxidant peroxidase | Redox homeostasis | A0A0H3MCJ5 |
| *thrS* | CTL0844 | 1.52 | 0.00501 | threonine-tRNA ligase | Translation | B0B8F5 |
| *aspS* | CTL0804 | 1.52 | 6.89E-03 | aspartate-tRNA ligase | Translation | B0B8B6 |
| *rpoD* | CTL0879 | 1.51 | 1.00E-02 | RNA polymerase sigma factor RpoD | Transcriptional regulation | A0A0H3MHM6 |
| *glyA* | CTL0691 | 1.47 | 0.00242 | serine hydroxymethyltransferase | Amino acid biosynthesis | B0B804 |
| *sctJ* | CTL0822 | 1.43 | 7.93E-03 | type III secretion protein | Type III Secretion | A0A0H3MDS0 |
| *rpoC* | CTL0566 | -1.28 | 7.87E-03 | DNA-directed RNA polymerase subunit beta' | Transcriptional regulation | B0B7N0 |
| *pGP8-D* | L2b_RS0475 | -1.36 | 4.91E-03 | virulence plasmid integrase pGP8-D | DNA replication and repair | B0BCM4 |
| *rplW* | CTL0788 | -1.41 | 0.0035 | ribosomal subunit | Translation | B0B8A0 |
| *prfA* | CTL0278 | -1.44 | 5.80E-03 | peptide chain release factor RF1 | Translation | B0B9D0 |
| *rplC* | CTL0790 | -1.52 | 4.30E-04 | ribosomal subunit | Translation | B0B8A2 |
| *CTL0061* | CTL0061 | -1.52 | 2.95E-03 | inorganic phosphate transporter PHO4 | Nutrient transport | A0A0H3MG71 |
| *CTL0659* | CTL0659 | -1.57 | 9.75E-04 | tetraacyldisaccharide 4'-kinase lpxK | Other | A0A0H3MC42 |
| *CTL0473* | CTL0473 | -1.57 | 1.22E-03 | hypothetical exported protein | Hypothetical | A0A0H3MBQ9 |
| *incD* | CTL0370 | -1.59 | 3.33E-03 | inclusion membrane protein D | Other | B0B9M3 |
| *plsX* | CTL0182 | -1.59 | 1.00E-02 | phosphate acyltransferase | Other | B0B939 |
| *CTL0613* | CTL0613 | -1.6 | 1.34E-03 | hypothetical inner membrane protein | Hypothetical | A0A0H3MC14 |
| *pmpA* | CTL0669 | -1.63 | 0.00534 | probable outer membrane protein PmpA | Other | A0A0H3ML49 |
| *CTL0548* | CTL0548 | -1.66 | 1.01E-03 | hypothetical non-heme Fe(II) 2-oxoglutarate | Hypothetical | A0A0H3MBY2 |
| *CTL0541* | CTL0541 | -1.67 | 1.00E-02 | hypothetical membrane protein | Hypothetical | A0A0H3MC35 |
| *sucB_2* | CTL0311 | -1.7 | 7.85E-03 | dihydrolipoyllysine-residue succinyltransferase | Energy Metabolism | A0A0H3ML42 |
| *amn* | CTL0120 | -1.71 | 4.71E-03 | AMP nucleosidase | DNA replication and repair | A0A0H3MGA3 |
| *mrsA* | CTL0547 | -1.77 | 1.00E-02 | phosphoglucomutase | Other | A0A0H3MC40 |
| *ftsY* | CTL0192 | -1.82 | 4.26E-04 | signal recognition particle receptor | Translation | A0A0H3MGE5 |
| *CTL0609* | CTL0609 | -1.85 | 7.29E-06 | hypothetical exported protein | Hypothetical | A0A0H3MDF7 |
| *dnaX_1* | CTL0439 | -1.92 | 5.15E-03 | DNA polymerase III subunit gamma/tau | DNA replication and repair | A0A0H3MBM8 |
| *CTL0314* | CTL0314 | -2.04 | 2.67E-04 | hypothetical membrane protein | Hypothetical | A0A0H3MGM7 |
| *CTL0430* | CTL0430 | -3.85 | 5.46E-05 | hypothetical integral membrane protein | Hypothetical | A0A0H3MBV0 |
| *CTL0063* | CTL0063 | -3.89 | 2.61E-03 | hypothetical | Hypothetical | A0A0H3MCG4 |

Up-regulation of *trpA* transcription during early-cycle iron-starvation was confirmed by RT-qPCR, while only modest increases were observed for the other up-regulated genes tested (Figure 6A). Down-regulation of *ctl0430*, *ctl0063*, and *incD* during iron-starvation could not be confirmed by RT-qPCR (Figure 6B). We reasoned that early-cycle responses were not detected by RT-qPCR for most of our tested genes due to the limit of detection of the technique. The raw values detected for most of our early-cycle transcripts fell at or below the lowest concentrations of our standard curves. Between 6 and 9h post-infection, chlamydial mRNA represents a very small proportion of the total RNA. This limitation was overcome for RNA-seq experiments by depleting rRNAs and eukaryotic RNA prior to synthesizing cDNA. However, cDNA used in RT-qPCR was prepared from total RNA because mRNA enrichment would have made it impossible for us to normalize our RT-qPCR data to chlamydial genomes. The overwhelming proportion of eukaryotic RNA present in the undiluted cDNA used as template may have impeded accurate detection of transcripts.

**Figure 6.**
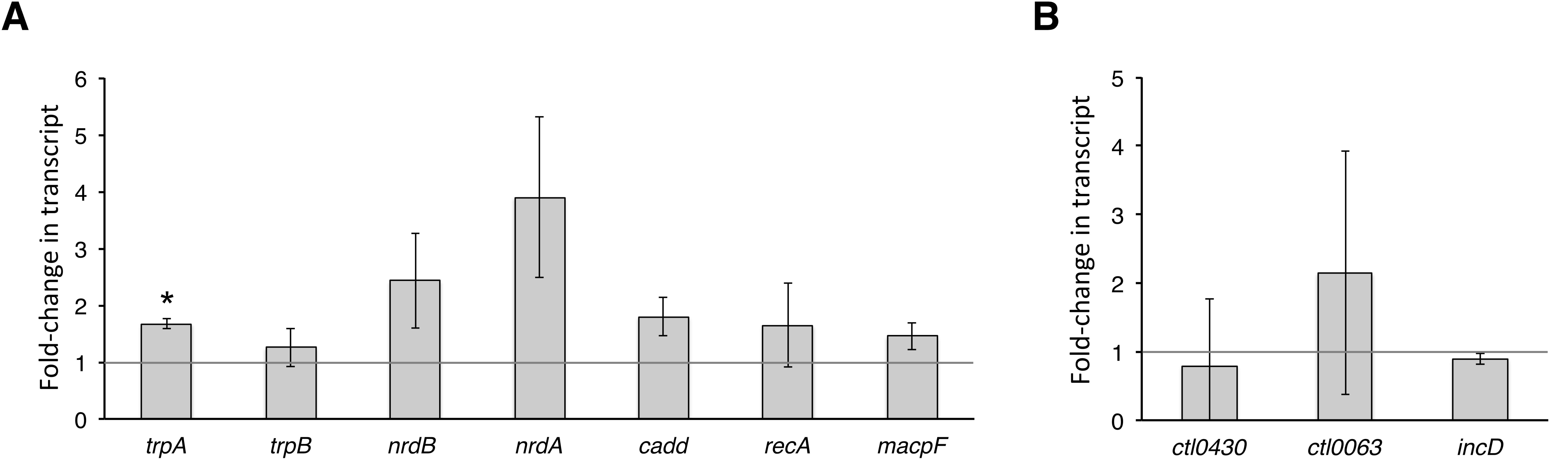
Confirmation of the early-cycle response to iron starvation by RT-qPCR. Transcripts that were significantly changed by RNA-sequencing, in response to iron-starvation starting at 6h p.i., were confirmed by RT-qPCR. A) Up-regulated or B) down-regulated transcription is indicated as fold-changes in transcripts after 3h BPDL-treatment in comparison to mock-treated at equivalent timepoints post-infection (solid gray bar). An asterisk indicates that the fold-change is significantly changed, with a p-value ≤ 0.05. Statistical analysis was done with a one-tailed Student’s t-test, based on two biological replicates.

### Network and biological pathway analysis

To further analyze the relevance of these gene expression changes to chlamydial survival, we utilized the bioinformatics tool STRING-db v.10.5 to generate networks of functionally associated genes (51). Representation of differentially expressed gene sets (p-value ≤ 0.05) from short-term iron starvation (3h) reveals gene networks with intersecting pathway clusters (manually added grey circles). We chose to use the less stringent p-value to allow entire pathways to emerge, which would not have been quite as apparent with a more stringent cutoff. Consistent with our predicted functional categories, network analysis of both early (Figure 7A) and mid-cycle (Figure 7B) responses to iron-starvation revealed clusters that include amino acid biosynthesis, DNA replication and repair, and translation. Functional clustering of the mid-cycle response also revealed the entire cluster of genes necessary to convert pyruvate to acetyl-CoA, as well as gene clusters involved in tRNA modification and charging.

**Figure 7.**
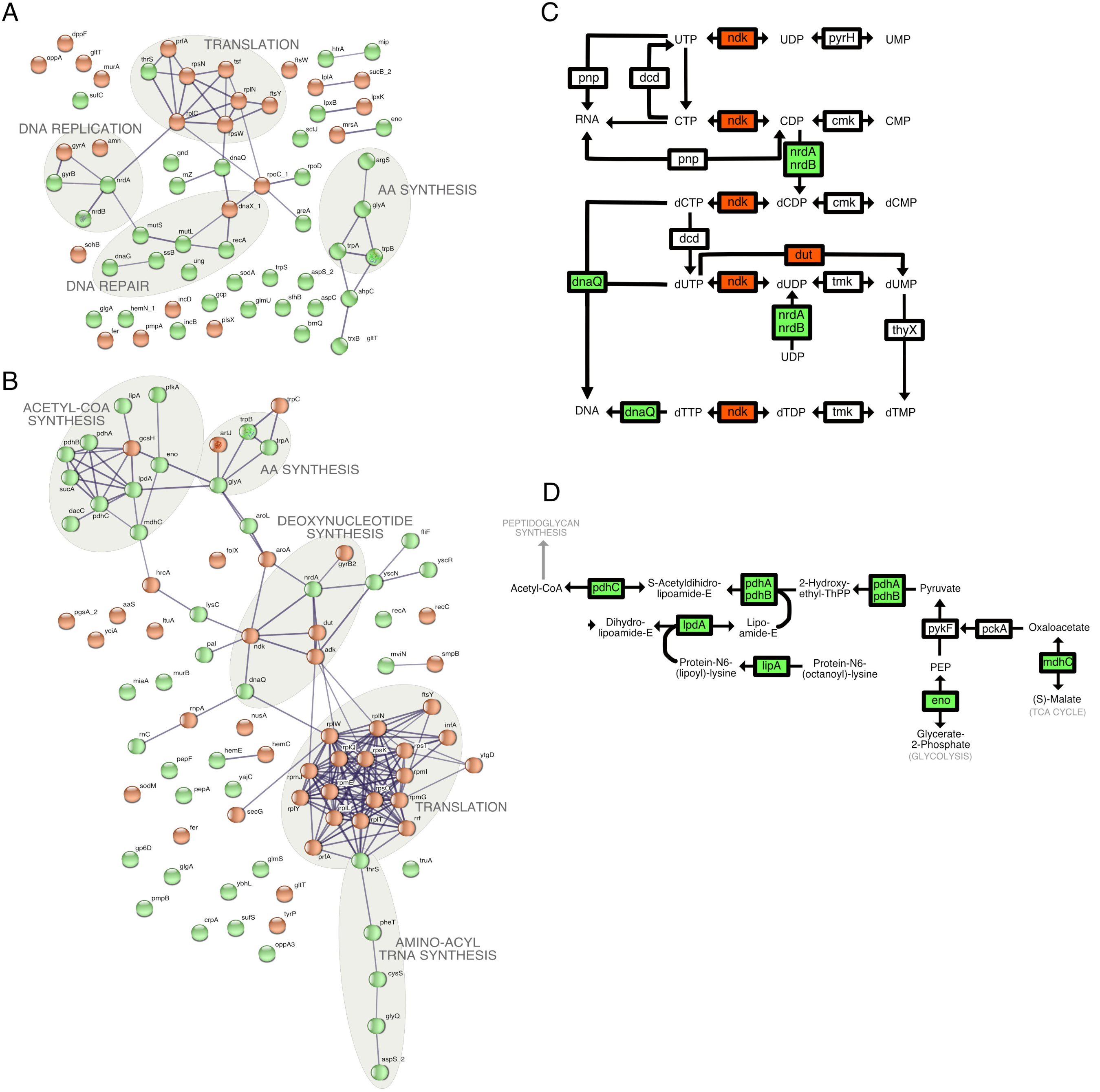
Pathway analysis of iron-starvation responses. Association networks for differentially expressed genes, with a p-value ≤ 0.05 and a minimum of 10 mapped reads were generated for the A) early-cycle response (6+3h BPDL) and B) mid-cycle response (12+3h BPDL) using STRING-db v.10.5. Line thickness connecting nodes (genes) correlates with confidence of gene association, with a minimum confidence cutoff of 0.7. Mid-cycle (12+3h BPDL or 12+6h BPDL) clustered genes were mapped to C) nucleotide metabolism and D) acetyl-CoA synthesis pathways using KEGGMapper v.2.8. Up-regulated and down-regulated genes in C) and D) are shown with green and red backgrounds, respectively. Unchanged genes have a white background.

The locus identifiers of genes in each identified cluster were submitted to KEGGMapper v2.8 to determine possible roles in specific biological pathways (52). For example, genes from the early-cycle DNA replication and repair cluster (Figure 7A) were mapped to multiple pathways including purine metabolism (5), pyrimidine metabolism (5), mismatch repair (5), replication (4), homologous recombination (3), double-stranded breaks repair (2), and base excision repair (1).

#### Nucleotide metabolism

We modified the KEGGMapper output for pyrimidine metabolism to indicate the direction of change in mid-cycle gene expression during iron starvation (Figure 7C). Under all iron-starvation conditions, the ribonucleotide reductase genes *nrdA* was up-regulated. Ribonucleotide diphosphates (NDPs) bound to NrdA are converted by NrdB to deoxynucleotide diphosphates (dNDPs). These dNDPs are not likely further converted to dNTPs, as indicated by the down-regulation of nucleotide diphosphate kinase, *ndk*. Available dUMP would likely be derived from the UDP pool, instead of from the dUTP pool, since transcription of the dUTP pyrophosphatase gene, *dut*, is down-regulated during iron-starvation. Taken together, these transcriptional changes would result in a net increase in dNDPs, enabling rapid DNA replication when iron levels and *ndk* expression return to normal (Figure 7C).

#### Amino acid biosynthesis

Functional clustering also indicates that *Chlamydia* prioritizes maintenance of amino acid pools during iron starvation. Multiple amino acid synthesis, inter-conversion, and uptake mechanisms were up-regulated in response to short-term iron-starvation. Transcriptional up-regulation of the branched chain amino acid transporter, *brnQ*, the aspartate aminotransferase, *aspC*, and the serine hydroxymethyltransferase, *glyA* may increase the diversity of the amino acid pool such that *Chlamydia* can quickly adapt to fluctuations in amino acids. Surprisingly, the tryptophan salvage pathway genes, *trpB* and *trpA*, were consistently up-regulated during short-term iron-starvation. The tryptophan synthase subunit TrpB catalyzes the beta-replacement of indole with serine to form tryptophan (Trp), while TrpA facilitates the interaction of TrpB with indole (53). Their role in recovery from IFN-g and Trp-starvation stresses is well documented, but differential regulation in response to iron-starvation is novel (54, 55). While the biological relevance of *trpBA* induction during iron-starvation is unclear, we reason that *Chlamydia* could in fact prepare for further immune insult (e.g. IFN-g induction of indoleamine 2,3-dioxygenase expression) by increasing intracellular Trp levels. Taken together, iron-starvation may increase levels of serine, aspartate, glutamate, branched-chain amino acids, and tryptophan, many of which are essential for normal development (56-60). Amino acid biosynthetic genes were significantly overrepresented (4.38-fold, p-value=0.0464) in the set of differentially expressed mid-cycle genes as determined by the Panther Overexpression Test (61).

#### Translation

The largest cluster generated from STRING-db included translation factors of the mid-cycle response (Figure 7B). Based on protein annotations in Uniprot and Biocyc databases, it is evident that *C. trachomatis* responds to iron-starvation by shutting down factors involved in every step of translation: ribosome assembly, initiation, elongation, termination, ribosome recycling, and peptide targeting (46, 54; Table 4). While preventing the assembly and function of translational machinery, *Chlamydia* also responds to iron starvation by increasing factors important for synthesis and modification of tRNAs, in addition to increasing transcription of *rnC*, the product of which cleaves rRNA transcripts into ribosomal subunit precursors (Table 4). Translation genes were significantly overrepresented (3.26-fold, p-value=0.0243) in the set of mid-cycle differentially expressed genes as determined by the Panther Overexpression Test (61).

**Table 4.**
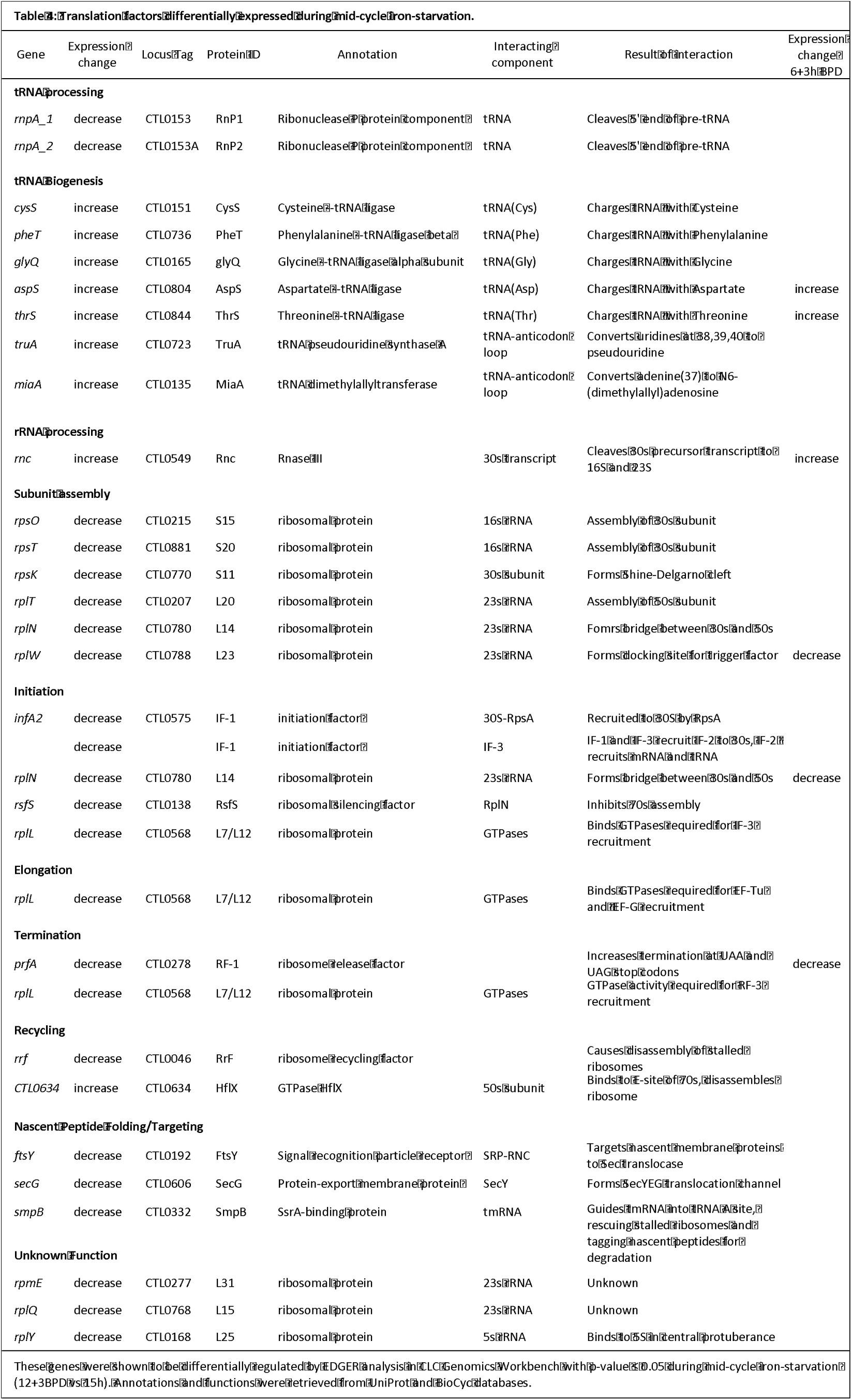

#### Acetyl-coA synthesis

Transcription of the entire set of genes necessary for conversion of pyruvate to acetyl-CoA was induced during the mid-cycle response to BPDL-treatment (Figure 7D). This includes the lipoylation enzymes *lipA* and *lpdA*, and the entire pyruvate dehydrogenase complex, *pdhABC*. In addition, transcription of the TCA cycle gene *mdhC* and glycolysis gene *eno* was induced, likely driving formation of pyruvate from different carbon sources. Acetyl-CoA can be converted to malonyl-CoA for fatty-acid biosynthesis or utilized in the formation of N-acetylglucosamine-1-phosphate for peptidoglycan synthesis, both of which are required for rapid growth of *Chlamydia* (63, 64). Since transcription of the peptidoglycan-modifying enzymes, *glmS* and *murB*, were also increased during iron starvation, acetyl-coA is likely used to form new peptidoglycan. Expression of fatty acid synthesis genes was unchanged during iron-starvation.

Pathway analysis of both early and mid-cycle responses to iron-starvation (Figure 7A and 7B) reveals that down-regulation of translation and up-regulation of amino acid synthesis and nucleotide synthesis may be important for surviving this stress. Similarly, a core set of 13 genes (*trpB, trpA, nrdA, recA, dnaQ, CTL0704, CTL0102, thrS, prfA, rplW, CTL0061, CTL0548, pGP8-D*) are differentially expressed after 3h BPDL treatment, during both early and mid-cycle responses, while *trpB, trpA*, and *nrdA* are up-regulated in all BPDL treatments (Fig S3). This overlap in differential gene expression is displayed as a Venn diagram in Supplemental Figure 3.

## DISCUSSION

We monitored the immediate global transcriptional response of *Chlamydia trachomatis* serovar L2 to short-term iron-starvation during early and mid-cycle (RB phase) development. In contrast to previous studies of iron starvation in *Chlamydia*, our short-term treatment with BPDL did not cause the hallmark changes in morphology and *euo* transcription associated with persistence. This approach enabled us to detect a response specific to iron starvation as *Chlamydia* tries to adapt to stress, rather than the transcriptome of the aberrant bacterium. By deep RNA-sequencing we were able to identify novel primary transcriptional responses, representing 7-8% of the genome, after only 3h of iron-starvation with BPDL. It is possible that a portion of the detected BPDL-responsive regulon is actually due to chelation of metals other than iron. Cu^2+^ is chelated at similar affinities to Fe^2+^ and Fe^3+^, while Zn^2+^ is chelated with 2-3 logs lower affinity than iron ions (65). We expect that Zn^2+^ is not efficiently depleted during the short-term BPDL treatments used in this study, but cannot exclude the possibility that we have detected transcriptional changes in response to altered availability of other metals. It is also possible that a more immediate response could be detected with even shorter BPDL treatments, though we expect a longer duration is required to chelate both free iron and iron bound to protein complexes in intracellular *Chlamydia*. Since only 12 genes were differentially expressed after 6h BPDL-treatment, a longer duration of treatment may be necessary to detect the full secondary response, which may not be obvious until the effects of the primary transcriptional response are realized at the protein level. This is supported by the fact that 6h BPDL-treatment maintains induction of the primary response operons, *trpBA* and *nrdAB*, while reducing or delaying expression of some late cycle genes (*scc2-ctl0840-copB-copD, tsp*). Decreased or delayed late gene expression has also been observed during long-term iron-starvation (43, 49, 66, 67).

In agreement with proteomic observations of deferoxamine-treated *C. trachomatis* after 24h and *C. pnuemoniae* after 48h post-infection, we observed up-regulation of *ctl0874*/*cadd*, *ahpC, eno, and htrA* during short-term BPDL-treatment (45, 68). In contrast to previous iron-starvation studies, we did not detect a significant increase in *ytgA* expression in our RNA-seq results. We expected the *ytgABCD* iron acquisition operon to be immediately induced in response to iron-starvation, since its repression by YtgR is dependent on available iron (42). Expression of the *ytgABCD* operon peaks during mid-cycle development, indicating that the iron-dependent repressor, YtgR, may be inactive or at low levels during early and mid-cycle (15, 42). It is possible that we do not observe significant differences in the expression of the operon during iron-starvation because it is already maximally expressed in the mock-treated controls (Table S3). Global detection of YtgR repression by CHIP-sequencing or targeted analysis of specific promoters will be necessary to delineate the contribution of YtgR activity to the detected iron-responsive regulon. Recent work to define targets of known transcription factors in *Waddlia chondophila* discovered binding-sites of YtgC by ChIP-sequencing (69). Interestingly, the most frequent target, *hrcA*, was also increased during iron-starvation in our study, indicating it may also be a target of YtgC in *C. trachomatis*.

Transcriptional responses to iron-starvation in most bacteria typically include up-regulation of iron-acquisition systems and virulence factors (70-72). While the *ytgABCD* operon was not up-regulated during short-term iron-starvation, other unidentified iron-uptake and iron-dependent repression mechanisms may exist, and thus could be represented in our set of iron-starvation induced genes. The virulence factor genes *cadd* (*ctl0874*) and *macpF* (*ctl0408*) were induced in both the early and mid-cycle response to iron-starvation. CADD overexpression induces apoptosis when expressed in cultured human epithelial cells, but has also been demonstrated to play a role in folate biosynthesis (73, 74). MACPF contains a domain that may enable perforin activity, but so far has only been show to undergo cleavage upon infection and become inserted into bacterial membranes (75). Several type III secretion structural components (*mcsC, sctJ, sctR, fliF, cdsN/ctl0043, cdsD/ctl0033*) and effectors (*ctl0884, ctl0476, ctl0184, ctl0081*) were also transcriptionally up-regulated during iron-starvation, which could potentially alter interactions between the host and chlamydial inclusion.

Similar to the up-regulation of the ribonucleotide reductase operon *nrdHIEF* during iron-starvation in *E. coli* and *Yersinia pestis*, the ribonucleotide reductase genes, *nrdA* and *nrdB*, are consistently up-regulated during under short-term iron starvation (71, 76). This up-regulation indicates that deoxynucleotides may be important for *Chlamydia* to survive this stress. However, since NrdB requires iron for its function, deoxynucleotide levels may not increase until iron becomes available. Instead, high levels of inactive NrdA-B complexes may actually impede replication and development by inducing stalling at replication forks, providing a possible explanation for the decreased replication observed during iron-starvation (77, 78).

*Chlamydia’s* immediate transcriptional response to iron-starvation is remarkably similar to stringent responses in other bacteria, which enable rapid adaptation to various stresses by diverting resources from macromolecular biosynthesis, e.g. translation, and growth to immediate survival, often resulting in a quiescent state (79, 80). This rapid transcriptional response is achieved through synthesis of the chemical alarmone, (p)ppGpp, which interacts with RNA polymerase and DksA to globally modify transcriptional activity (81, 82). During amino acid starvation in bacteria, uncharged tRNAs in the A-site of ribosomes are sensed by RelA, which responds by synthesizing (p)ppGpp from ATP and GDP or GTP (83, 84). (p)ppGpp can also be synthesized and hydrolyzed by SpoT during other stress conditions. However, since *Chlamydia* lacks the RelA and SpoT homologues necessary for (p)ppGpp synthesis, they likely evolved alternative mechanisms to reduce growth and increase survival responses during stress (17, 85, 86). Iron-starvation has been shown to induce a stringent response in *Bacillus subtilis* that up-regulates transcription of amino acid biosynthesis genes (87).

Multiple amino acid synthesis, inter-conversion, and uptake mechanisms were up-regulated in response to short-term iron starvation. Surprisingly, the primary response includes an increase in transcripts involved in tryptophan salvage, *trpB* and *trpA*, but not the tryptophan-dependent repressor *trpR*. TrpR-dependent regulation of the polycistronic transcript, *trpRBA*, has been extensively studied during tryptophan starvation and IFN-g treatment, but rarely, if ever, in the context of iron-starvation (54, 88, 89). Notably, *trpB*, but not *trpR* levels were also increased under estradiol-induced persistence, suggesting a *trpR*-independent mechanism of inducing tryptophan salvage transcription may exist (90).

Pathway analysis clearly indicates that transcripts involved in all steps of translation from initiation to ribosome recycling are down-regulated during iron-starvation. This reduction in translation factors may lead to an eventual shutdown or modification of translation activity that could increase survival during stress. By shutting down energy expensive protein synthesis, ATP and GTP pools can be rerouted to immediate survival responses (tRNA-charging, transcription). Similarly, iron-starvation reduces the transcription of several ABC transporter genes, which require ATP for their function. Uncoupled RNA and protein levels in *Chlamydia* have also been observed during IFN-g stress (17). The apparent decrease in translation during IFN-gamma exposure could be exacerbated by decreases in the levels of components of the translation machinery in response to simultaneous iron-starvation. However, decreased expression of translation factors during the primary response to iron-starvation may not be apparent until preexisting ribosomal-protein complexes are degraded or destabilized. This may explain why ≥24h of iron-starvation is required to induce the development of aberrant RBs (43). Down-regulation of translation factors during iron starvation will have to be examined at the protein level to determine its contribution to adaptation to iron starvation and development of persistence.

In contrast to down-regulation of translation, iron starvation increases transcription of amino-acyl synthesis genes (*cysS, pheT, glyQ, aspS, thrS*), which are responsible for charging tRNAs with amino acids. The apparent disconnect between increased amino-acyl-tRNA pools and decreased translation indicates possible survival mechanisms. Charged tRNAs might be utilized in an immediate survival response to iron-starvation, prior to the turnover of ribosomal subunits. Alternatively, *Chlamydia* may be accumulating charged tRNAs for recovery and resumption of development when normal levels of iron and translation factors are restored.

A major theme that emerged from our gene expression analysis is that *Chlamydia* likely perceives iron starvation as a signal to prepare for further nutrient deprivation and immune insult. Transcriptional up-regulation of tryptophan salvage pathway (trpB, trpA), oxidative stress (*ahpC*, *pdi*, and *sodM*) and DNA repair (*mutS, mutL, ssb, ung, recA*) genes indicate a protective response against antimicrobial insults of the inflammatory immune response (e.g. IDO activation, reactive oxygen species). As an obligate intracellular pathogen, *Chlamydia* has undergone reductive evolution with constant selective pressure from the host immune system and its multiple anti-chlamydial effectors. Due to its small genome (~ 1 Mbp), *Chlamydia* may not have the capability to induce a specific transcriptional response to each particular stressor, and the simultaneous deployment of stress responses may have been the most parsimonious route of adaptation to immune insult. In this case, we would expect that iron-starved *Chlamydia* would be better protected from damage by antimicrobial insults than mock-treated *Chlamydia*. Immediate transcriptional responses to other stress conditions will need to be monitored to determine if this coordination of antimicrobial responses is unique to iron starvation.

This study provides the first evidence of a global iron-dependent regulon for *C. trachomatis*. By using a systems-approach to delineate *Chlamydia*’s transcriptional response to iron starvation, we have been able to detect biological pathways and place them in the context of chlamydial development. These findings are novel, and add to previous studies of iron-dependent transcriptional and proteomic profiling in aberrant RBs, revealing transcriptional adaptive strategies prior to the development of a persistent state. Additionally, our results include a high-resolution profile of mid-cycle development of *C. trachomatis* serovar L2, including relevant time points for monitoring shifts in early, mid, and late-cycle gene expression. We expect this dataset will prove useful for future studies that seek to determine *Chlamydia*’s immediate transcriptional response to other chemical and/or nutrient stresses. Our findings include previously unrecognized shifts in energy utilization and down-regulation of translation that resemble a stringent-like survival response. *Chlamydia* may utilize a two-stage approach of increasing transcription of survival genes in the short term to delay development and survive during iron-starvation, followed by an eventual shutdown of translation at later times of sustained stress. The latter might account for the observed irreversibility of the persistent state during long-term starvation for iron or tryptophan.

## MATERIALS AND METHODS

### Cell Culture and infection

HeLa monolayers were infected with *C. trachomatis* strain L2 434/Bu in 6-well plates at a multiplicity of infection (MOI) of 2 for RNA and genomic DNA (gDNA) collection experiments, and on coverslips in 24-well plates for morphology studies. Cells were grown in DMEM supplemented with 10% FBS, 2 mM glutamine, and 10 micrograms/mL gentamycin in 5% CO_2_ at 37°C. HeLa cells used in this study were started from P1 stocks from ATCC, and were regularly checked for contamination by DAPI-staining and the Universal Mycoplasma detection kit (ATCC).

### RNA-sequencing

RNA was collected and pooled from 2 or 4 T75 flasks of *C. trachomatis*-infected HeLa monolayers that had been treated with 100 μM 2,2-Bipyridyl (BPDL) starting at 6 or 12h hours post-infection (6h + 3h BPDL, 12h + 3h BPDL, 12h + 6h BPDL) and in mock-treated samples at equivalent timepoints post-infection (9h, 12h, 15h, 18h). RNA was purified using the RiboPure Bacteria (Ambion) kit as per manufacturers instructions. Total RNA was further enriched for transcripts over 100 nucleotides in length with the MegaClear kit (Ambion). Mammalian transcripts and rRNAs were removed with the MicrobEnrich kit (Ambion) and bacterial rRNAs were removed using the MicrobExpress kit (Ambion), repeating 2-3 times. The integrity and quantity of total and depleted RNA was monitored with an AATI Fragment Analyzer. cDNA libraries were prepared with the Ion Total RNA-seq Kit V2, and sequencing beads prepared using an ion Chef, and sequencing performed on an Ion Proton chip with HiQ chemistry. Primary sequence analysis, trimming, and binning of reads was performed using Torrent Suite Software version 5.0.5. Remaining reads were mapped to the combined core genome of *C. trachomatis* strain L2 434/Bu (Genbank Accession: AM884176) and the plasmid of *C. trachomatis* L2b CS784/08 (NZ_CP009926) using CLC Genomics Workbench 9, requiring reads be at least 30 nucleotides in length, with default alignment parameters.

The EdgeR algorithm in CLC Genomics was used to determine differential gene expression during development and iron-starvation, assuming a false-discovery rate of 10% and p-values ≤ 0.05. tRNAs and ribosomal RNAs were filtered from the reads to account for differences in depletion efficiency, and only genes with at least 5 mapped reads were included in the analysis. Differentially expressed genes were confirmed for selected transcripts by RT-qPCR.

### qPCR and RT-qPCR

*C. trachomatis*-infected HeLa monolayers were treated with 100 micromolar BPDL starting at 6 or 12h hours post-infection (6h + 3h BPDL, 12h + 3h BPDL, 12h + 6h BPDL) and mock-treated samples at equivalent timepoints post-infection (6h, 9h, 12h, 15h, 18h). RNA and gDNA were collected with the RiboPure Bacteria and the DNeasy Blood and Tissue (Qiagen) kits, respectively. cDNA was generated with Superscript IV reverse transcriptase (Life Technologies) from 200-500 nanograms RNA as per manufacturer’s instruction, except the use of random nonamers instead of hexamers. Transcripts were amplified with PowerUp SYBR Green system from undiluted cDNA for early (6-9h) or diluted 1:10 in 10 mM Tris for midcycle (12-18h) samples and detected with Applied Biosystems 7300 RT-qPCR system.

### Chlamydial morphology

Chlamydiae were monitored for changes in morphology in response to mock-treatment or treatment with 100 micromolar BPDL starting at 12h post-infection for 3, 6, or 12h. Infected cultures were fixed on coverslips and stained with pooled human serum (Sigma, H4522) at 1:750, followed by goat-anti-human antibody conjugated to Alexa Fluor 488 (ThermoFisher) at 1:1000. DNA was stained with DAPI at 5 micrograms/mL. Images were taken on a Leica SP8 confocal microscope with a 63X oil-objective and 4X zoom.

### IFU assay

Chlamydiae were monitored for changes in infectivity in response to mock-treatment or treatment with 100 micromolar BPDL starting at 12h post-infection for 12 or 24h, at an MOI of 1. Infected cultures were scraped into 300 microliters SPG and stored at −80 C for later testing. Thawed lysates were serially diluted into complete DMEM, centrifuged onto HeLa monolayers in 24-well plates, washed with HBSS, and allowed to infect for 24h. Infected cultures were fixed and stained with pooled human serum at 1:750, followed by goat-anti-human antibody conjugated to Alexa Fluor 488 (ThermoFisher) at 1:1000. Inclusions were counted by fluorescent microscopy and inclusion-forming units (IFU) were calculated as previously described.

### Visual analysis of differentially expressed genes

Functional categories were assigned for all genes differentially regulated with a p-value ≤ 0.01 by referring to the GO terms listed on UniProt. Pie charts were generated using the “pie” function in Rstudio. Heatmaps were generated in Rstudio using the package “pheatmaps”, with parameters set to average clustering and Euclidean distance. The Panther overexpression test in Panther v12.0 was done on differentially regulated gene sets (total) with p-values ≤ 0.01, using the default parameters and Bonferri correction. Pathway analysis was performed on differentially regulated genesets with p-values ≤ 0.05 and a minimum of 10 mapped reads, with STRING-db v.10.5 set to confidence ≥ 0.7. StringDB maps were slightly modified to make space to increase font size, indicate the direction of change by color-coding and add pathway labels, without altering network relationships. Clustered genes detected with StringDB were further analyzed using KeggMapper v.2.8, and pathway maps were generated based on KeggMapper output using Affinity Designer v1.4.1.

## Data availability

Raw and processed sequencing files were submitted to the NCBI Gene Expression Omnibus (GEO) as a Superseries, and the mid-cycle and early-cycle projects can be found using accession number GSE106763.

## ACKNOWLEGEMENTS

We would like to acknowledge Dr. Scot Ouellette and Nicholas Pokorzynski for critical reading of the manuscript. This work was funded by start-up funds for R.A.C. from the School of Molecular Biosciences, College of Veterinary Medicine at Washington State University, NIH grant 2R01AI065545-06A1 awarded to R.A.C., and NIH training grant T32AI7025-36 for A.J.B.

## SUPPLEMENTARY DATA

**Figure S1. Annotated heatmap of BPDL-treated and mock-treated gene expression in *C. trachomatis* that corresponds to Figure 2A.**

**Figure S2. Annotated heatmap of mid-cycle iron-starvation corresponding to the subset in** **Figure 2B**.

**Figure S3. Venn diagram of differential gene expression for all BPDL-treatments**. Overlap in genes that were differentially up-regulated or down-regulated across multiple treatments are displayed as a Venn diagram (p-value ≤ 0.01).

**Table S1. Summary of RNA-sequencing and mapping in this study**.

**Table S2. Normalized means of mock-treated and BPDL-treated transcription during mid-cycle development of *C. trachomatis.*** The mean and log_10_ mean expression values of genes that showed a significant change in gene expression during normal mid-cycle development, p-value ≤ 0.01 are displayed for the following EdgeR comparisons: 12h vs 18h, 12h vs 15h, 15h vs 18h) These values were used to create the heatmaps in Figure 2A. Genes that had at least one value that was greater than the 4.5 threshold have an asterisk, and the values are displayed in bold.

**Table S3. Complete expression profile of *C. trachomatis* during normal development**. The RNA-sequencing reads and EdgeR analysis of normal development of *C. trachomatis* was exported from CLC Genomics Workbench 9.5.3. Samples were normalized across the entire dataset by quantile scaling. rRNAs, tRNAs, and Features (genes) with less than 10 reads in all samples were eliminated from the dataset prior to normalization and EDGE analysis. Unique Reads are raw values. Samples were merged from multiple chips to obtain a minimum of 8X coverage.

**Table S4. Complete expression profile of *C. trachomatis* during mid-cycle iron starvation**. The RNA-sequencing reads and EdgeR analysis of iron-starved *C. trachomatis* was exported from CLC Genomics Workbench 9.5.3. Samples were normalized across the entire dataset by quantile scaling. rRNAs, tRNAs, and Features (genes) with less than 10 reads in all samples were eliminated from the dataset prior to normalization and EDGE analysis. Unique Reads are raw values. Samples were merged from multiple RNA-sequencing chips to obtain a minimum of 8X coverage.

**Table S5. Complete expression profile of *C. trachomatis* during early-cycle iron starvation**. The RNA-sequencing reads and analysis of iron-starved *C. trachomatis* was exported from CLC Genomics Workbench 9.5.3. Samples were normalized across the entire dataset by quantile scaling. rRNAs, tRNAs, and Features (genes) with less than 10 reads in all samples were eliminated from the dataset prior to normalization and EDGE analysis. Unique Reads are raw values. Samples were merged from multiple RNA-sequencing chips to obtain a minimum of 8X coverage.

**Table S6. Primers used in this study**.

